# The Dynamics of *Cryptococcus neoformans* infection in *Galleria mellonella*

**DOI:** 10.1101/2025.03.19.644221

**Authors:** Daniel F. Q. Smith, Aviv Bergman, Arturo Casadevall

## Abstract

*Galleria mellonella* has emerged as an important host for the study of fungal virulence, insect immune responses, and the evaluation of antifungal agents. In this study we investigated the dynamics of fungal infections in *G. mellonella* using *Cryptococcus neoformans*, a human pathogenic fungus. Since the analysis of infection dynamics requires a fine temporal resolution of larval death, we employed a photographic timelapse technique that allowed us to simultaneously measure death by proxy of larval melanization and absence of movement. Larval mortality occurred in two phases, early and late, which differed in their timing of melanization. Early phase deaths occurred with rapid whole-body onset of melanization, followed by sudden cessation of movement several hours later. Contrastingly, late phase deaths occurred with a gradual cessation of movement, followed by melanization, typically radiating from one location on the larva. The differences in mortality kinetics suggests differences in fungal pathogenesis with one population succumbing early while the rest linger for later death. Subsequent analysis of mortality data using the inversion method revealed predictable deterministic dynamics without evidence for chaotic signatures, indicating that this *C. neoformans-G. mellonella* infection model behaves differently than bacterial-insect models.

**Importance:** The ability to predict the course of an infection is critical to anticipating disease progression and effectively treating patients. Similarly, the ability to make predictions about pathogenesis in laboratory infection models could further our understanding of pathogenesis and lead to new treatments. As fungal diseases are expected to rise, understanding the dynamics of fungal infections will be important to anticipate and mitigate future threats. Here, we developed a timelapse method to visualize infections of *Galleria mellonella* larvae with the fungal pathogen *Cryptococcus neoformans*. This method provided insight into infection progression that are not apparent from standard survival measurement protocols, including the relationship between melanization and death. Further, it enabled us to explore the dynamics of disease progression in this system, which revealed deterministic dynamics without evidence of chaos, implying predictability in the outcome of cryptococcal infection in this moth.

## Introduction

Interactions between hosts and microbes vary greatly between host species, microbial species, and specific circumstances of the interaction. Even when comparing the outcome of interactions of the same host and microbial species in different experiments or situations, changes in experimental conditions (i.e. microbial inoculum, temperature, or state of the hosts’ immune systems) can have major effects on pathogenic potential, symptoms of disease, and the outcome of infection (1, 2). A host infected with a microbe at a certain inoculum might be able to clear the infection readily, while a host exposed to the same inoculum but with a slightly different immune response might launch a host-damaging inflammatory response to the microbe, or might fail to clear the infection, resulting in microbial growth and host death. Hence, damage can come from both the host immune response and microbial action (1). Understanding the sources of variability in the outcome of host-microbe interactions is important for improving reproducibility, anticipating future threats and developing improved therapies for infectious diseases.

A fundamental question in the field of infectious diseases is the predictability of host-microbe interactions. To know whether the host-pathogen interactions variables are predictable, one must first determine whether the dynamics of the system, which can be either random (stochastic) or non-random (deterministic). The outcome in random dynamics is not predictable. For non-random or deterministic dynamical systems, one must further differentiate whether it is chaotic (non-predictable) or non-chaotic (predictable). Predictability would facilitate understanding and treatment of infections, because given certain parameters, clinicians could effectively predict the trajectory of pathogenesis and mitigate it. If the host-microbe interactions are chaotic, it means that while there are non-random variables at play, the system is too dependent on the initial conditions, which implies non-predictability. Mathematical chaos is typified in popular culture by the metaphor of the “butterfly effect,” when seemingly insignificant actions or variables (i.e. a butterfly flying), can lead to unpredictable and significant changes in larger and distant systems (I.e. altered weather patterns). While chaotic systems, such as weather, are amenable to short term predication, the countless minuscule, yet inevitably significant variables, cause downstream effects that long term prediction impossible. If a host-microbial system is chaotic in a manner like weather systems, understanding the important inputs of to the chaotic host-microbe interactions can allow the creation of prediction models in a manner similar to modern meteorology. Chaos also affects the reproducibility of experiments, a timely topic given concerns about the reproducibility of results in the biomedical sciences (3).

A prior study of host-microbe dynamics used bacterial infection of invertebrate model hosts (4). That study found evidence of chaos in the lifespan of insects and helminths infected with *Pseudomonas* spp., but not in the lifespan of control uninfected organisms. While bacteria are important pathogens that affect hosts across kingdoms of life, that study did not include fungal infections. Using different methodology and low sample size, *C. neoformans* infection in *Galleria mellonella* wax moth larvae was found to be deterministic without evidence for chaos (5). Comparing bacterial and fungal infections is important because fungi exhibit different strategies for pathogenesis and differ from bacteria in their dependence between their pathogenic potential and inoculum. Bacteria, which largely cause disease through the release of toxins into the host, kill hosts in a direct inoculum-dependent manner according to measures of pathogenic potential (6). Contrastingly, fungi, which cause disease largely through survival and growth in the host, do not kill in a directly dose-dependent manner; rather, there is a logarithmic relationship where the order of magnitude of inoculum leads to higher death and measures of pathogenic potential (6). This implies different strategies for bacterial and fungal pathogenesis that might be reflected in the dynamics of their interaction

A widely used system for the study fungal pathogenesis is the *Galleria mellonella* wax moth model (7–12). *G. mellonella* allow for easy screening of virulence of different fungal isolates or mutants, in a high-throughput and relatively rapid, easy, and affordable manner due to fast infection timelines and easy availability of the organism. Here we used C. neoformans infection in *G. mellonella* to explore the dynamics of animal fungal infection. We developed a timelapse imaging protocol to track *G. mellonella* larval survival at a much higher resolution of time intervals than is traditionally used (15 minutes versus 24 h). Using this system we found no evidence for chaotic signatures in this model.

## Results/Discussion

There are three current limitations with the collection of survival data in the *G. mellonella* model for the purpose of studying infection dynamics. The first is the lack of temporal resolution for the death event, since this is usually measured when the experimenter checks the larvae. and it is not logistically feasible to achieve near continuous monitoring. The second is that death is determined by manually checking the movement of the larvae at each time interval, which usually involves physical stimulation of the larvae with a pipette tip, which stresses the animal and could affect the results. Third is the need to collect hundreds of survival data points for sufficient statistical power needed for the chaos calculations. To help overcome these limitations, we developed a survival monitoring model that allowed frequent survival measurements and removed the necessity of manual probing to ascertain whether the animal was alive or dead. Specifically, we assembled timelapse cameras to record the movement of *G. mellonella* larvae following infection within 24-well plates, with one larva per well. This allowed each larva to be individually monitored at increments of 15 minutes. We analyzed the timelapse movie frames and recorded the time at which movements of the larvae ceased as well as the time in which melanization began to spread through the larvae’s bodies. Larval melanization results from the production of melanin pigment, which is part of the insect immune response, and is associated with imminent or recent larval death (13–15).

The time lapse photography setup allowed us to study the kinetics of infected larva melanization relative to the cessation of movement (Supplementary Video 1). We first compared cessation of movement as measured by time lapse photography to the accepted method of manual larval poking for establishing death (Figure 1A). The photography and manual poking survival curves closely paralleled one another but there was an approximate 1-day lag in mortality as measured by cessation of movement, implying that some of immobile larvae could be roused to move if poked. The fact that cessation of movement was highly correlated with eventual demise we accepted this difference in mortality timing because it eliminated mechanical poking, which could potentially introduce a set of new variables ranging from stress to operator variability and renders high temporal resolution almost impossible. We experimented with placing larvae in 12- and 24-well plates. The attraction of the 24-well plate was that it would allow us to image a larger number of larvae per experiment. However, larvae placed in 24-well plates died before those in 12-well plates (Figure 1B). Although this phenomenon was not further investigated it could reflect additional stress from confinement in a small space. Alternatively, reducing space would reduce their space for movement, which is necessary for hemolymph circulation. regardless, we settled on using 24-well plates even though this reduced our experimental survival time.

**Figure 1.**
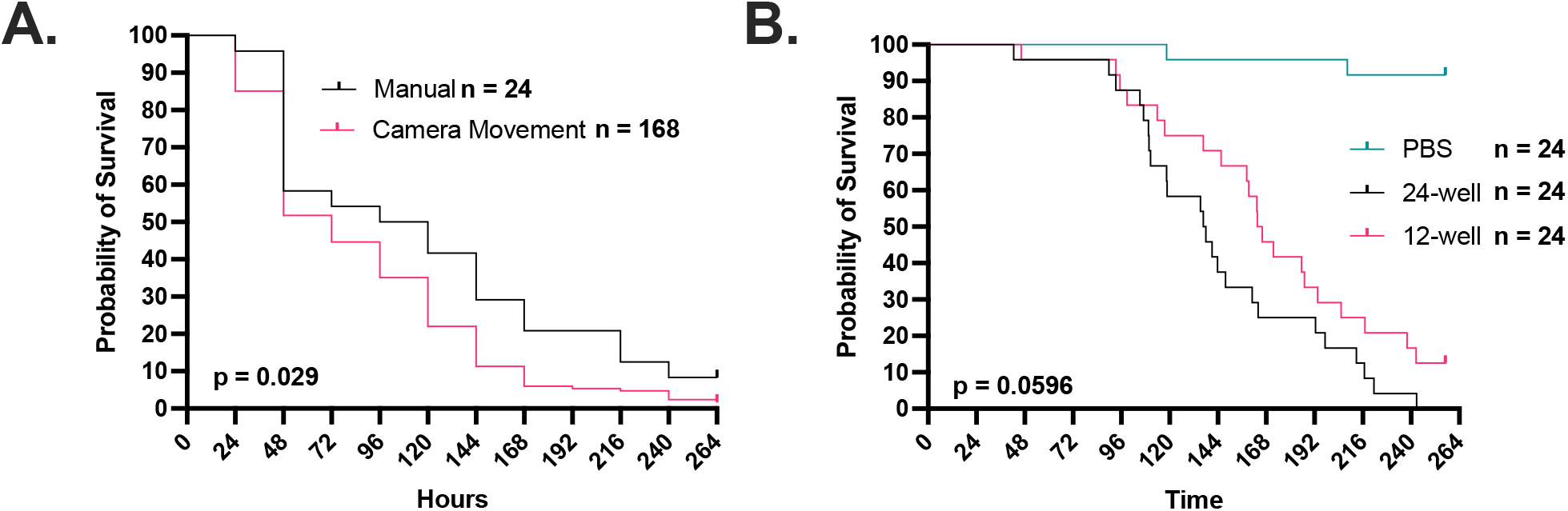
Timelapse imaging can be used to record G. mellonella movement as a proxy for survival. **A**. The use of cessation of movement and time of melanization as a proxy for larval death is consistent with the gold standard of manually checking the larvae every 24 h with a physical stimulus. Camera movement survival data was rounded to nearest 24 h increment. Manual survival group has an n = 24, while the group with survival quantified from the camera timelapse movement has n = 168. **B**. Survival of larvae in a 12-well plate is improved compared to those kept in a 24-well plate. All groups in Panel B have n = 24. P-values represent log-rank Mantel-Cox comparisons

Next, we used time lapse photography to evaluate the dynamics of cryptococcal infection in this host. *G. mellonella* survival after a high inoculum infection with *C. neoformans* manifested two distinct phases, one concluding by 48 h (Supplementary Video 2) and the other starting gradually at about 96 h (Supplementary Video 3) and continuing till the end of the experiment when all the larvae died (Figure 2A). Melanization preceded the cessation of movement during the first phase generally followed the cessation of movement during the second later phase (Figure 2A). With control larvae that died following PBS injection, which presumably reflect trauma, death in these controls occurred later than the deaths from *C. neoformans* infection, and the time between melanization and cessation of movement was closely linked. When the same survival data were rounded to the nearest 24-h increment, as would be the case with the standard daily manual survival measurements, we see that this two-phase curve is lost due to the reduced temporal resolution (Figure 2B). This indicates that some patterns of host survival and clues to underlying pathogenesis are usually lost due to standard survival assay methodology based on recording events at discrete time intervals.

**Figure 2.**
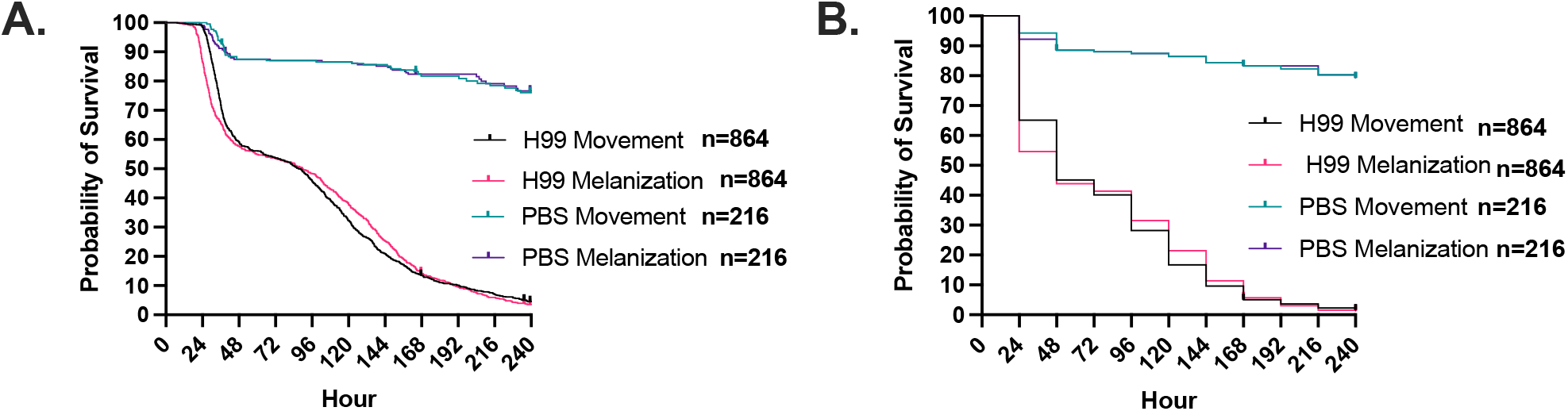
*C. neoformans* infection of *G. mellonella* at high temporal resolution shows two phases of infection. **A**. Using timelapse photography, *G. mellonella* larval survival following infection with *C. neoformans* was monitored in increments of 15 minutes, and time cessation of movement and onset of melanization was recorded for each larva. This showed a two-phase survival curve with one large phase of death occurring at 48 h and another slower phase after 96 h. **B**. The appearance of these two phases is lost when the temporal resolution is lowered to every 24 h. For H99 infections, n = 864, while the PBS-injected n = 216.

Melanization in *G. mellonella* larvae was measured by quantifying the mean gray value of the individual wells containing the larvae. Melanization causes the larvae to become darker and thus the mean gray value of the well becomes lower since lower gray values denote darker pixels. When looking at the melanization response we noticed that early phase larvae melanized rapidly and uniformly throughout their bodies, approximately 5-10 h prior to the cessation of movement (Figure 3A, B, C, E). Larval melanin reached a brief plateau immediately prior to the cessation of movement (Figure 3C, red arrows), followed by more melanization. Conversely, the larvae that died later had melanization start in one spot of their body and then spread (Figure 3A, D, E). This spot was usually either at the head or the posterior end, in a location in the body cavity not necessarily associated with the site of injection. The melanization reaction then spread slowly from these spots and/or from the head of the larvae for several hours after the larvae stopped moving (Figure 3E). The white arrow indicates where melanization began and radiated from, in addition to the head of the organism (Figure 3E). This again suggests two mechanisms of pathogenesis during infection. The rapid and complete onset of melanization in the early death phase indicates a systemic melanization response in the hemolymph of the larvae. Since melanin synthesis is the result of a polymerization reaction that yields highly reactive and toxic intermediates, melanization prior to death suggests the possibility that this immune mechanism is contributing to the demise of the larvae. In contrast, the focal melanization in the later phase could reflect melanization within a large immune nodule (16) – a complex of immune cells, clotting factors, melanization factors, and other immune components aimed at restricting and killing microbes. One interpretation of these results is that early deaths occur when the fungus is dispersed throughout the hemolymph triggering widespread melanization, while later deaths reflect following dissemination from an infected tissue or an immune nodule that is no longer able to contain the fungal infection.

**Figure 3.**
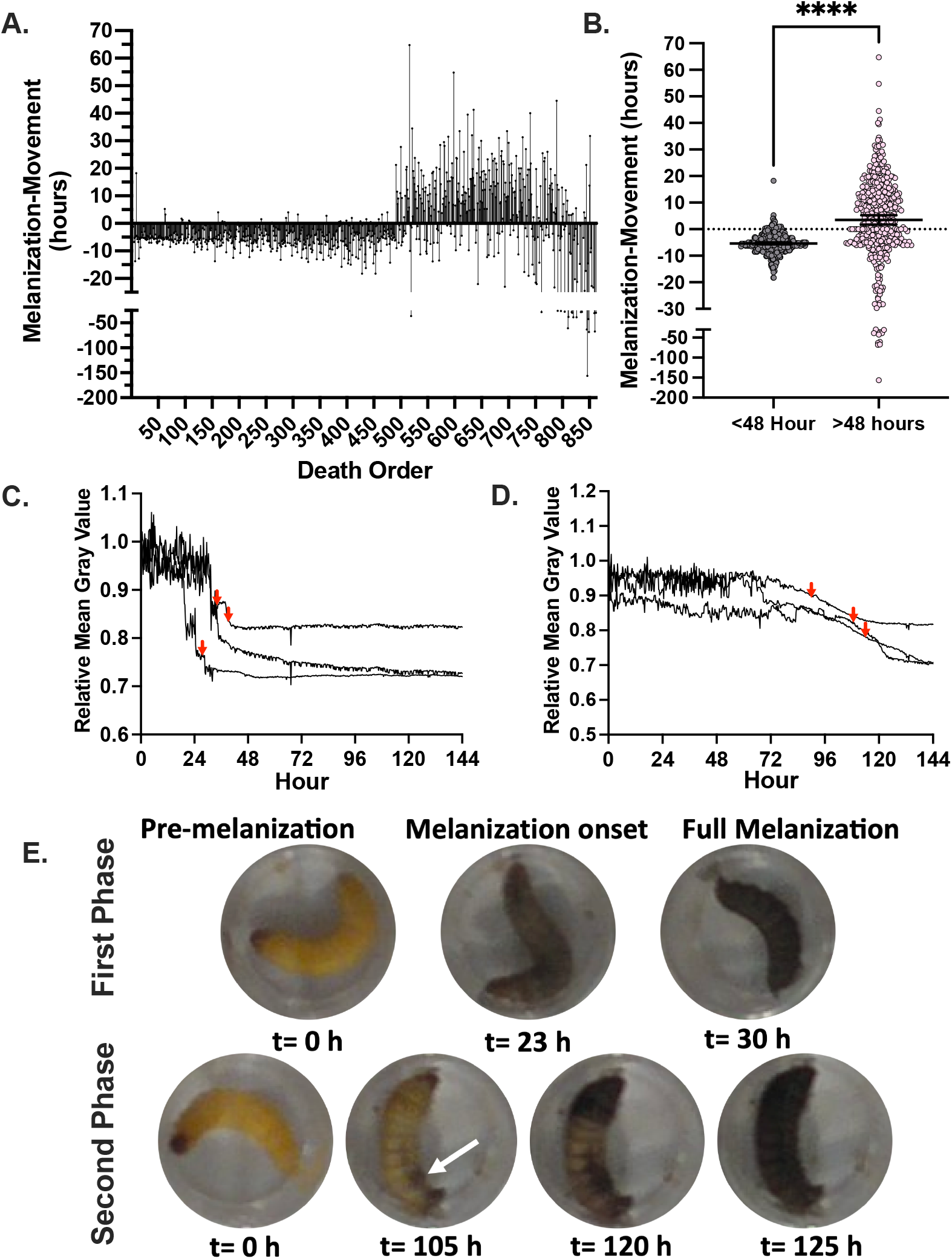
*G. mellonella* larvae show di8erent characteristic melanization responses upon infection. **A**. Larvae that die during the first phase of death show onset of the melanization response approximately 5 h prior to the larvae stop moving, while those at later time points have more mixed patterns of melanization, often with pigmentation occurring hours after the larvae stop moving. **B**. The diRerence between melanization and cessation of movement in the first 48 h (n = 459) is diRerent than the value for deaths occurring after 48 h (n=405). Statistical significance was determined through an unpaired t-test, **** represents p<0.0001. **C**. Larvae from the first phase of death show a sharp decline in mean gray value, which indicates rapid production of black melanin pigment, **(D)** while larvae from the second phase of death show a later and more gradual reduction in the mean gray value. **Panels C and D** are representative quantifications of mean gray value for 3 separate larvae. Red arrows indicate time at which the respective larvae stopped moving. **E**. Representative images of a larva that died in the first phase of death compared to a larva from the second phase of death. Note that for early phase there is continued larval movement following the onset of melanization, while the larva in the second death phase does not move following melanization onset. White arrow indicates spot in which melanization began from.

To examine the dynamics of the *C. neoformans-G. mellonella* interaction, we applied the inversion measure to the observed time distributions. This approach allowed us to assess deviations from expected stochastic patterns by comparing our empirical data to a null hypothesis of non-chaotic behavior. Our analysis did not yield statistically significant evidence to reject the null hypothesis (*p* ≥ 0.2) (Figure 4A), suggesting that, within the resolution of our dataset and analytical framework, there is no strong indication of chaos governing the observed host-pathogen interactions. Given that our observations were taken at 15 min, deaths recorded at a given time point must have reflected events occurring within that time interval. To account for this impreciseness in knowing the timing of the event, we randomly assigned observed deaths across smaller subintervals within that interval, introducing randomness into the analysis. While this approximates the likely times of death, it could also obscure any chaotic signature. Consequently, rather than calculating a single inversion measure for the histogram, we generated a distribution of inversion measures and compared it to the null bootstrap distribution. No statistically significant difference was found between the sample and the bootstrap null distributions (*p* = 0.5) (Figure 4B). This results are consistent with a prior analysis of the dynamics of *C. neoformans-G. mellonella* interactions using different analytic methods that also found no chaos in this system (5).

**Figure 4.**
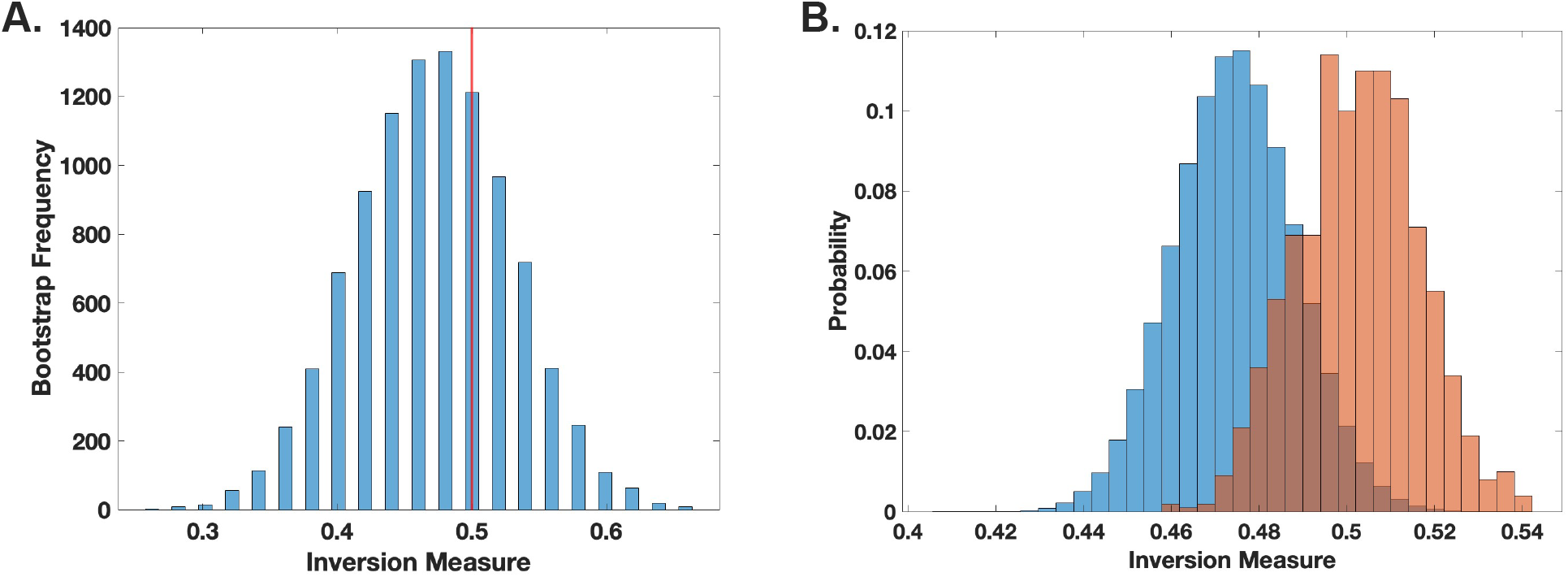
Death of G. mellonella larvae following C. neoformans infection does not demonstrate chaotic signatures. **(A)**. The inversion method using bootstrap for locally linear approximation of the distributions (histograms) was applied. The red line represents the actual inversion measure of the distribution. No statistical significance was found, indicating no clear evidence of chaotic behavior. (**B)**. Histogram comparing the null (blue) and sample (orange-red) distributions of the inversion measure. Inversion measure was performed through analysis of 842 larval death events.

The absence of chaotic signatures in this system differs from evidence of chaos in *Pseudomonas* spp. infections of flies and worms (4). We do not know whether this reflects a fundamental difference between bacterial and fungal infections, is a consequence of using a different host, a different method of infection or another aspect of the experimental inoculation. Since bacteria differ from fungi in mostly kill their host in a linear and logarithmic inoculum-dependent manner, respectively (6), it is possible that the difference in dynamics reflects an intrinsic difference between these microbial types. An additional difference between this study and the prior bacterial studies is that those infections were acquired through natural inoculation through ingestion by the host. In contrast, in this study we injected larvae with the inoculum directly since there is no practical means of inducing a natural cryptococcal infection in *G. mellonella*. Hence, our approach involved piercing the larval surface with a needed and delivering the inoculum to deeper tissues, which could produce a non-chaotic deterministic outcome by causing a fulminant infection that abrogates chaotic signatures. Future studies will have to dissect these possibilities to determine whether fungal infections show signs of mathematical chaos or whether the reason for the lack of chaos in these findings is a result of experimental design and infection procedures. Those studies will need to be done with other fungal pathogens, such as those entomopathogenic fungi that have co-evolved with insects and are thus able to naturally infect the larvae without the need for injection or other forced traumatic inoculation.

Another possibility is that a chaotic signature does exist in this system, but the results are a false negative and its detection was hindered by limitations in the available data and the method used. Chaotic dynamics can be highly sensitive to initial conditions and may require a sufficiently large and high-resolution dataset to capture subtle fluctuations that signify chaotic behavior. If the dataset is too sparse, contains significant observational gaps, or lacks a sufficiently long time series, any underlying chaotic patterns may remain undetected or appear as weak signals obscured by noise. Further data collection with higher temporal resolution and greater replication may help clarify whether chaotic signatures are present but undetectable under current conditions. Additionally, the inverse method employed for chaotic detection may require refinement or revision to improve its accuracy in this specific biological context. Different methods for identifying chaos, such as Lyapunov exponent analysis, recurrence quantification analysis, or state-space reconstruction, may vary in their sensitivity to noise and sampling limitations. Hence, it is possible that the analytic approach used here does not adequately capture the nonlinear dynamics characteristic of fungal infections, particularly if fungal-host interactions involve more complex regulatory feedback mechanisms than bacterial infections. Future studies could explore alternative or complementary analytical techniques, refining the methodological framework to better discern chaotic signatures in host-fungal interactions.

An additional consideration when judging the implications of these findings for other systems is our use of *G. mellonella*. Virulence in *G. mellonella* models of mutants or different clinical isolates of the fungus *C. neoformans* corresponds to virulence in standard murine models (8). While the *G. mellonella* model is good for assessing virulence through survival and fungal burden assays, it lacks an adaptive immune response, which may lead to differences with mammalian infection where the adaptive immune response is important. As of now, there are no readily available inbred strains for this insect. While this reflects the situation for infection in outbred populations it can also lead to variability between infected larvae depending on the genetic differences between individuals. Phenotypic and genetic variability in hosts may lead to noise in outcome measurements, resulting in chaotic signatures signal loss.

In summary, we developed a new method to study cryptococcal infection in *G. mellonella* based on timelapse recording of the movement and pigmentation dynamics of individual larvae, which provides a way to assess infection dynamics and disease progression at high temporal resolution and in a high-throughput environment. Analysis of the mortality outcomes from cryptococcal infection yielded deterministic non-chaotic dynamics with the caveat that we cannot rule out the existence of non-apparent chaotic dynamics.

## Methods

### Galleria mellonella Infections

Final instar *Galleria mellonella* larvae were obtained from Vanderhoorst Wholesale Inc. (St. Mary’s, OH, USA). Larvae were left to acclimate overnight in weighing boats. Larvae were then injected with 10^6^ cells of *C. neoformans* strain H99 suspended in PBS into the left rear proleg using a 1 mL insulin needle 28 ½ gauge and Stepper Injector. The average volume delivered to each larvae was 10 µl. Larvae were then placed into 24-well plates and kept at room temperature.

### Survival Recordings

Timelapse imaging of the infected larvae was performed at room temperature over 10 days. Pairs of 12- or 24-well plates were imaged with a Brinno TLC130 Time Lapse Camera, positioned from above using a clamp stand with the capture rate set to once every 15 minutes (Supplementary Figure 1). Following the completion of the experiment, the timelapse images were transferred and viewed using FIJI (ImageJ) (17). Control survival determinations, where survival was determined by movement following physical stimulus with a pipette tip, were performed concurrently.

### Movement Analysis

Timelapse movies were viewed on FIJI (ImageJ) so that each frame could be analyzed individually (17). We manually scanned through the movement of each larva and recorded frame after which no further movement was observed. This was recorded as the time at which movement stopped. To obtain hour of death, the frame number was divided by four.

### Melanization Analysis and Quantification

Timelapse movies were processed on FIJI (ImageJ) (17) with each frame analyzed individually. We scanned through each frame of the timelapse and recorded the frame in which melanization was first seen occurring in the larvae. This was recorded as the time that melanization occurred.

To further quantify the dynamics of melanization within the larvae, each of the wells in the 24-well plate were selected using the circular selection tool in FIJI and added as different Regions of Interest (ROIs) using the ROI Manager tool. The multi measure tool was selected, which then measured the mean gray value of each well during each of the timelapse frames. From this, we can see a sharp drop in mean gray value associated with the onset and acceleration of melanization within the larvae.

### The inversion measure on a time distribution

Given a distribution of time points, we first construct a histogram. For naturally discrete time points, as in our simulated waiting times derived from chaotic or stochastic processes, each unique time point serves as a bin. For continuous processes, we partition the distribution into *n* bins based on a chosen parameter *n*.

Given a histogram with n bins, each containing an integer count, we first introduce a small uniform random noise between 0 and *ϵ* (*ϵ* < 1) to break ties while preserving the relative order of distinct counts. Next, we partition the bins into consecutive, non-overlapping groups of four, discarding any remainder. For each sequence *x*_1_, *x*_2_, *x*_3_, *x*_4_, we define a **countertrend** (or **inversion**) as occurring when (*x*_4_ - *x*_1_) and (*x*_3_ - *x*_2_) share the same sign (both positive or both negative). We then compute the proportion of sequences exhibiting inversions. Since this frequency is influenced by the randomness of the tie-breaking step, we repeat the process 1000 times with different randomizations and report the average.

To compute a p-value for the inversion measure, we performed a bootstrap test against the null hypothesis that the histogram is smooth. We considered two types of null densities: **kernel-smoothed** and **locally linear**.

For kernel smoothing, we applied MATLAB’s built-in ksdensity function to the sample histogram. For the locally linear null density, we linearized the histogram as follows: for each consecutive sequence of four bins, let *x*_1_, *x*_2_, *x*_3_, *x*_4_ represent the corresponding whole-number values. We fit a line of best fit for *x*_i_ as a function of ii and use this line as the null density for that sequence. If any fitted values fall below zero, we reset them to zero. Repeating this process for all sequences of four bins results in a piecewise linear null density, which is then normalized to sum to one.

To generate a bootstrap sample from a null density (either kernel-smoothed or locally linear), we draw the same number of samples as in the original histogram and recalculate the inversion measure. This procedure is repeated 1,000 times to create a bootstrapped null distribution of the inversion measure. We then computed a t-statistic and its corresponding p-value by comparing the observed inversion measure to this null distribution.

For the *Galleria* datasets, where some observations were not recorded at regular intervals, we addressed gaps by redistributing events as follows: whenever one or more observations were missing, we uniformly distributed the next observed count of events across the interval between the most recent and prior observations. A histogram was then constructed from this redistributed data, and the inversion measure was computed as before.

Since the redistribution process introduces randomness, we repeated it 1,000 times, generating a sample distribution of inversion measures. For each of these 1,000 histograms, we conducted a bootstrap procedure with a sample size of 1,000, ultimately producing a null distribution composed of 1,000,000 inversion measures. Using these distributions, we estimated the *p*-value by calculating the probability that the sample inversion measure was greater than or equal to a value drawn from the null distribution.

## Data availability

Survival data used in this analysis and mean gray value measurements are deposited on FigShare under the DOI:10.6084/m9.figshare.28616303

## Acknowledgements

D.F.Q.S. is supported by NIH T32 AI 0074170-29. A.C. and D.F.Q.S are supported in part by R01 AI152078, R01 HL059842, and AI052733. A.B. was supported in part by NIH R01-CA164468 and R01-DA033788. Biorender was used to produce Supplementary Figure 1.

## Figure Legends

**Supplementary Figure 1.**
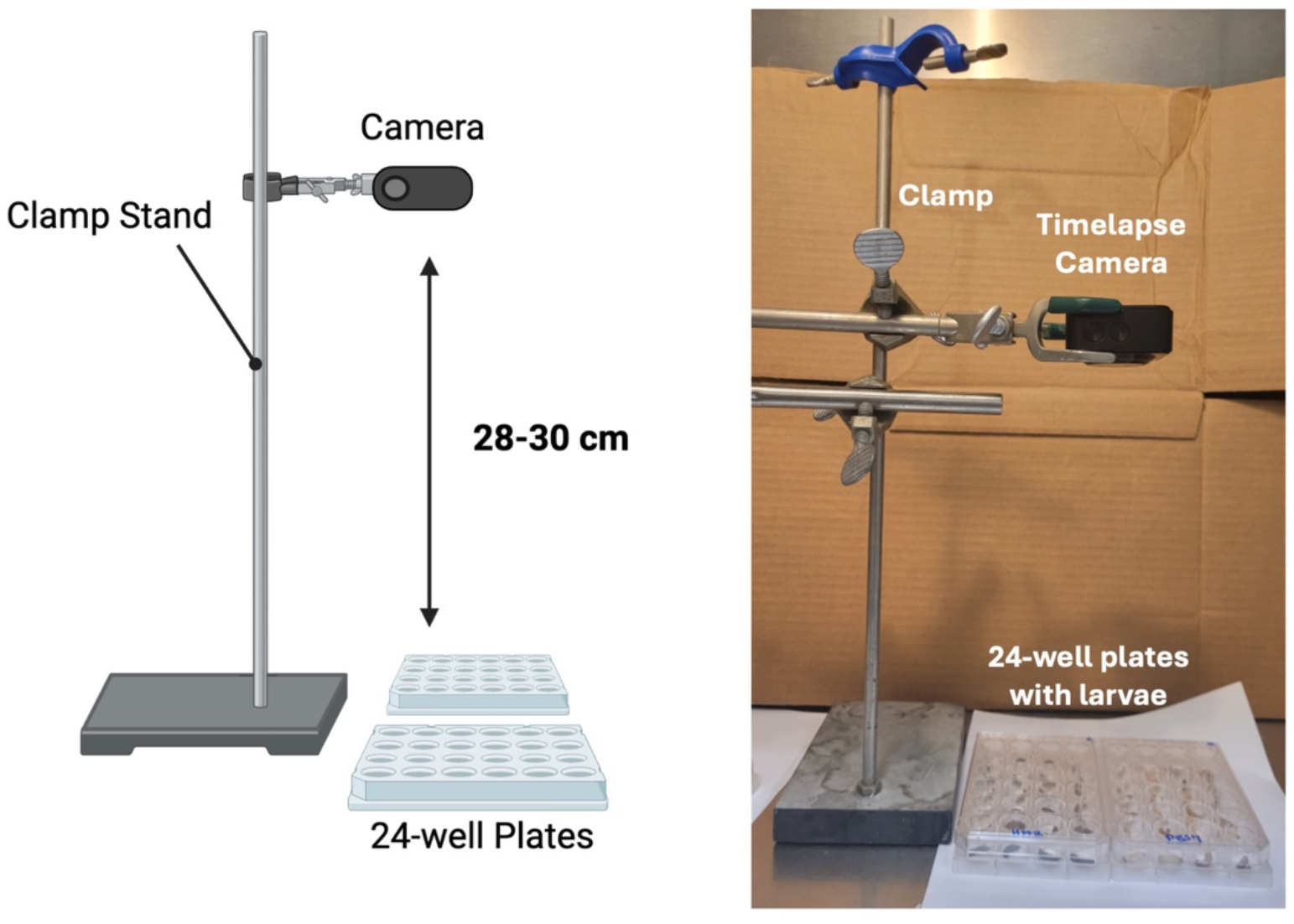
Timelapse photography set up. Timelapse camera is suspended above two 24-well plates containing *G. mellonella* larvae using a 3-pronged clamp attached to a support base.

